# Insight into the genomic history of the Near East from whole-genome sequences and genotypes of Yemenis

**DOI:** 10.1101/749341

**Authors:** Marc Haber, Riyadh Saif-Ali, Molham Al-Habori, Yuan Chen, Daniel E. Platt, Chris Tyler-Smith, Yali Xue

## Abstract

We report high-coverage whole-genome sequencing data from 46 Yemeni individuals as well as genome-wide genotyping data from 169 Yemenis from diverse locations. We use this dataset to define the genetic diversity in Yemen and how it relates to people elsewhere in the Near East. Yemen is a vast region with substantial cultural and geographic diversity, but we found little genetic structure correlating with geography among the Yemenis – probably reflecting continuous movement of people between the regions. African ancestry from admixture in the past 800 years is widespread in Yemen and is the main contributor to the country’s limited genetic structure, with some individuals in Hudayda and Hadramout having up to 20% of their genetic ancestry from Africa. In contrast, individuals from Maarib appear to have been genetically isolated from the African gene flow and thus have genomes likely to reflect Yemen’s ancestry before the admixture. This ancestry was comparable to the ancestry present during the Bronze Age in the distant Northern regions of the Near East. After the Bronze Age, the South and North of the Near East therefore followed different genetic trajectories: in the North the Levantines admixed with a Eurasian population carrying steppe ancestry whose impact never reached as far south as the Yemen, where people instead admixed with Africans leading to the genetic structure observed in the Near East today.

## Text

Yemen’s location and history suggest it has played an important role in early human migrations and thus is of considerable importance for understanding modern human genetic diversity.^1^ But Yemen as well as the rest of the Near East have so far been poorly represented in databases that catalogue human variation: The 1000 Genomes Project did not include any Near Eastern samples among its many populations from around the globe,^2^ and the Human Genome Diversity Project included only samples from the Northern region of the Near East.^3^ Here, we continue our efforts^4; 5^ to catalogue the diversity of genetically under-represented populations by genotyping and whole-genome sequencing individuals from Yemen, and provide the new data as a resource for future genetic studies in the region. We present an initial exploration of the data by asking how present-day Near Easterners are related to each other and to the ancient people who inhabited the Near East before them, focusing on the historical events and processes that have contributed to the genetic structure in Yemen today.

Samples were recruited from diverse locations in Yemen (Figure 1, Table 1) and DNA extraction from 3 ml blood performed in Yemen at Sanaa University. All subjects provided an informed consent form and the study was approved by the IRB of the Lebanese American University. Samples (n=169) were genotyped at the Wellcome Sanger Institute in the UK using the Illumina HumanOmni2.5-8 BeadChip which covers 2.5 million SNPs. In addition, 46 samples were whole-genome sequenced at >30× depth using the Illumina HiSeq X Ten and the reads were mapped to build 37 and processed as described previously.^6^ The use of the samples in genetic studies was approved by the Human Materials and Data Management Committee at the Wellcome Sanger Institute (HMDMC approval number 14/072).

**Table 1.** Newly genotyped and sequenced individuals and their location in Yemen.

| Location in Yemen | Genotyped Individuals | Sequenced Individuals | Latitude | Longitude |
| --- | --- | --- | --- | --- |
| Abyan | 6 | 4 | 13.568675 | 46.132999 |
| Amran | 6 | 4 | 15.66155 | 43.946018 |
| Bayda | 3 | 3 | 14.326473 | 45.380295 |
| Dali | 8 | 0 | 13.922642 | 44.786675 |
| Dhamar | 6 | 4 | 14.542997 | 44.400061 |
| Hadramout | 20 | 4 | 15.55487 | 50.418282 |
| Hajjah | 8 | 0 | 16.117894 | 43.03823 |
| Hudayda | 35 | 5 | 14.797868 | 42.954487 |
| Ibb | 17 | 4 | 14.02017 | 44.103111 |
| Jawf | 10 | 0 | 16.613066 | 45.616588 |
| Lahij | 5 | 2 | 13.05581 | 44.88286 |
| Mahwit | 3 | 4 | 15.365839 | 43.460814 |
| Maarib | 4 | 4 | 15.416376 | 45.35 |
| Saada | 5 | 4 | 16.950941 | 43.747772 |
| Sanaa | 10 | 0 | 15.36198 | 44.201962 |
| Shabwa | 9 | 0 | 15.108985 | 46.72996 |
| Taizz | 14 | 4 | 13.575224 | 44.021527 |
| **Total** | **169** | **46** |  |  |

**Figure 1.**
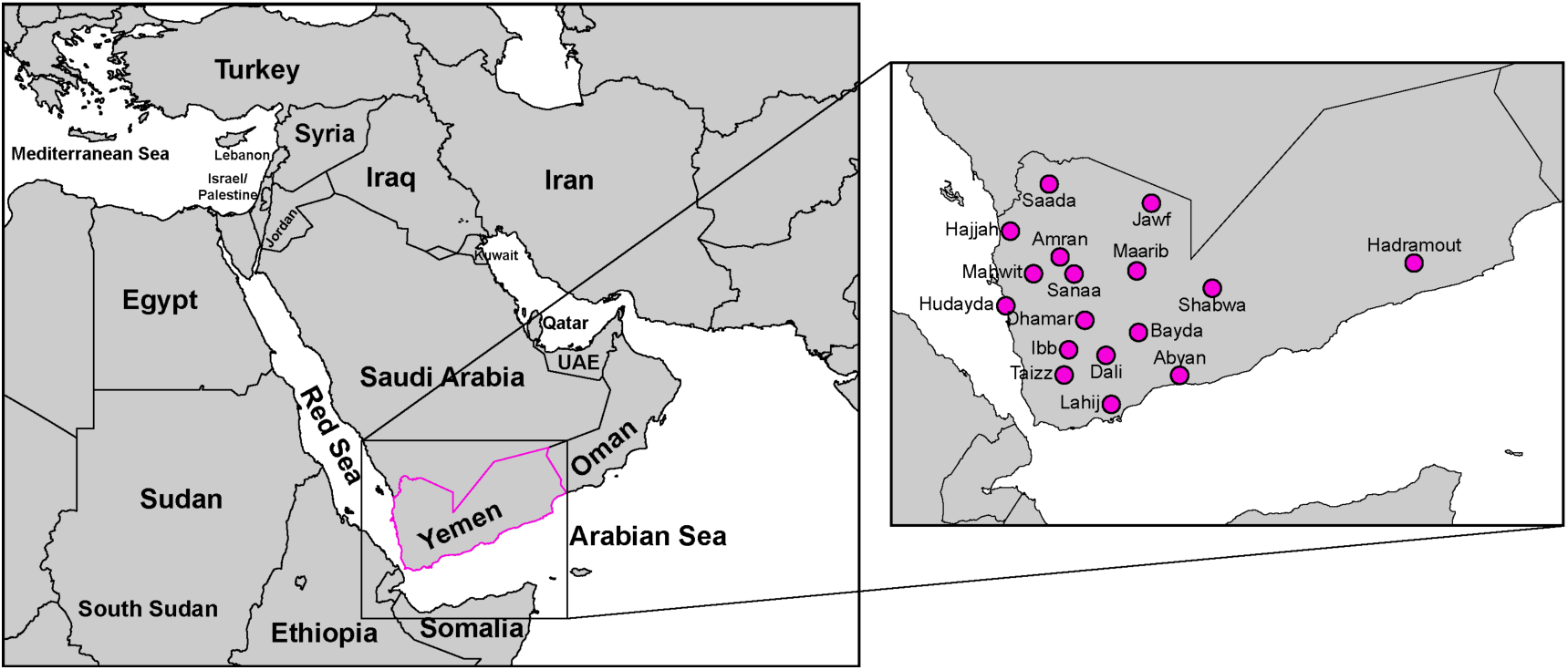
Map of the Near East with inset showing the location of the samples studied in this work.

We merged the newly generated data with previously published data from modern and ancient populations creating two datasets for the analyses: 1) The *geno set* included the new Yemen genotype data merged with the Human Origin dataset^7-9^ and with published ancient DNA data^4; 7; 8; 10-13^ resulting in a set of 3060 samples and 211,966 SNPs. 2) The *seq* set included the new Yemen sequencing data merged with data from the Simons Genome Diversity Project^14^ and with sequencing data from Lebanon^4^ in addition to published ancient DNA data,^4; 7; 8; 10-13^ resulting in a set of 731 individuals and 938,848 SNPs. In this work we refer to Yemen as being the South of the Near East, while Lebanon and the surrounding Levant regions are considered the North of the Near East. This differentiation is based on geography but also captures our previous knowledge of the genetic structure in the region.^5^

The newly sequenced data contained 1,212,228 SNPs not previously reported in dbSNP build 151. Among the novel SNPs we found 46,970 SNPs that were common (>5%) in the Yemeni population; annotation on build 37 using VEP v79 as described previously^15^ showed these included 336 moderately common missense SNPs, 23 splice donor/acceptor SNPs and 10 stop gained SNPs at >2% frequency (Table 2) which could be relevant to future medical studies in the region.

**Table 2.** Novel variants discovered in Yemen; their type and frequency.*

| Type | >0--<=2% | >2--<=5% | >5--<=50% |
| --- | --- | --- | --- |
| 3_prime_UTR_variant | 5643 | 1488 | 194 |
| 5_prime_UTR_variant | 1142 | 161 | 33 |
| TF_binding_site_variant | 242 | 93 | 24 |
| intergenic_variant | 296000 | 164117 | 19253 |
| intron_variant | 433468 | 220645 | 25712 |
| mature_miRNA_variant | 12 | 6 | 1 |
| missense_variant | 2352 | 259 | 77 |
| non_coding_transcript_exon_variant | 4855 | 1553 | 248 |
| regulatory_region_variant | 22554 | 8502 | 1358 |
| splice_acceptor_variant | 50 | 13 | 1 |
| splice_donor_variant | 46 | 7 | 2 |
| splice_region_variant | 519 | 81 | 26 |
| start_lost | 6 | 2 | 0 |
| stop_gained | 69 | 9 | 1 |
| stop_lost | 3 | 0 | 0 |
| synonymous_variant | 1196 | 165 | 40 |

\*Annotations are from VEP v79 using build 37.

Investigation of the demographic history of the Yemeni populations by applying multiple sequentially Markovian coalescent (MSMC)^16^ analyses to the Yemen genomes showed a typical Eurasian population size change pattern characterized by a bottleneck around 50,000-60,000 years, followed by a population size expansion (Figure 2). But the Yemen population size increase after the bottleneck appears to be limited compared with the increase which occurred in the North of the Near East (Figure 3). To understand how the Yemenis were related to other populations in the Near East, we projected them and the ancient individuals in the *geno* set onto the first two principal components computed from the variations in present-day West Eurasians and East Africans. The PCA (Figure 4 and Figure 5) revealed genetic structure between the North and South of the Near East, with most of the ancient Neolithic and Bronze Age Levantines clustering at intermediate positions between the two regions. We observe on the PCA two genetic clines that affect the Near Easterners differently: 1) The African cline, which appears to be a major contributor to genetic diversity in Yemen and 2) The Eurasian cline, towards which the Northern Near Easterners appear shifted compared with the Yemenis. The Northern Near Easterners are themselves structured on the African cline with Palestinians, Jordanians, Syrians, and Lebanese Muslims having more African ancestry than Assyrians, Jews, Druze and Lebanese Christians (Figure 4) confirming our previous observation.^5^ We investigated the time when admixture has occurred in North and South of the Near East by using linkage disequilibrium,^17; 18^ using the Lebanese Muslims and the Yemenis to represent the two areas, and setting Africans, East Asians and West Eurasians in the *geno* set as references (Figure 6). We found two significant admixture events in the Lebanese Muslims; the first occurred around 600BCE-500CE (Z=3.5) and the second occurred around 1580CE-1750CE (Z=3.4), confirming our previous results on the date of admixture in North of the Near East.^19^ In contrast, we detect one significant admixture event in Yemen occurring 1190CE-1290CE (Z=14.6) and thus these results suggest that the shifts of the North and South of the Near East along the African cline could arise from independent events.

**Figure 2.**
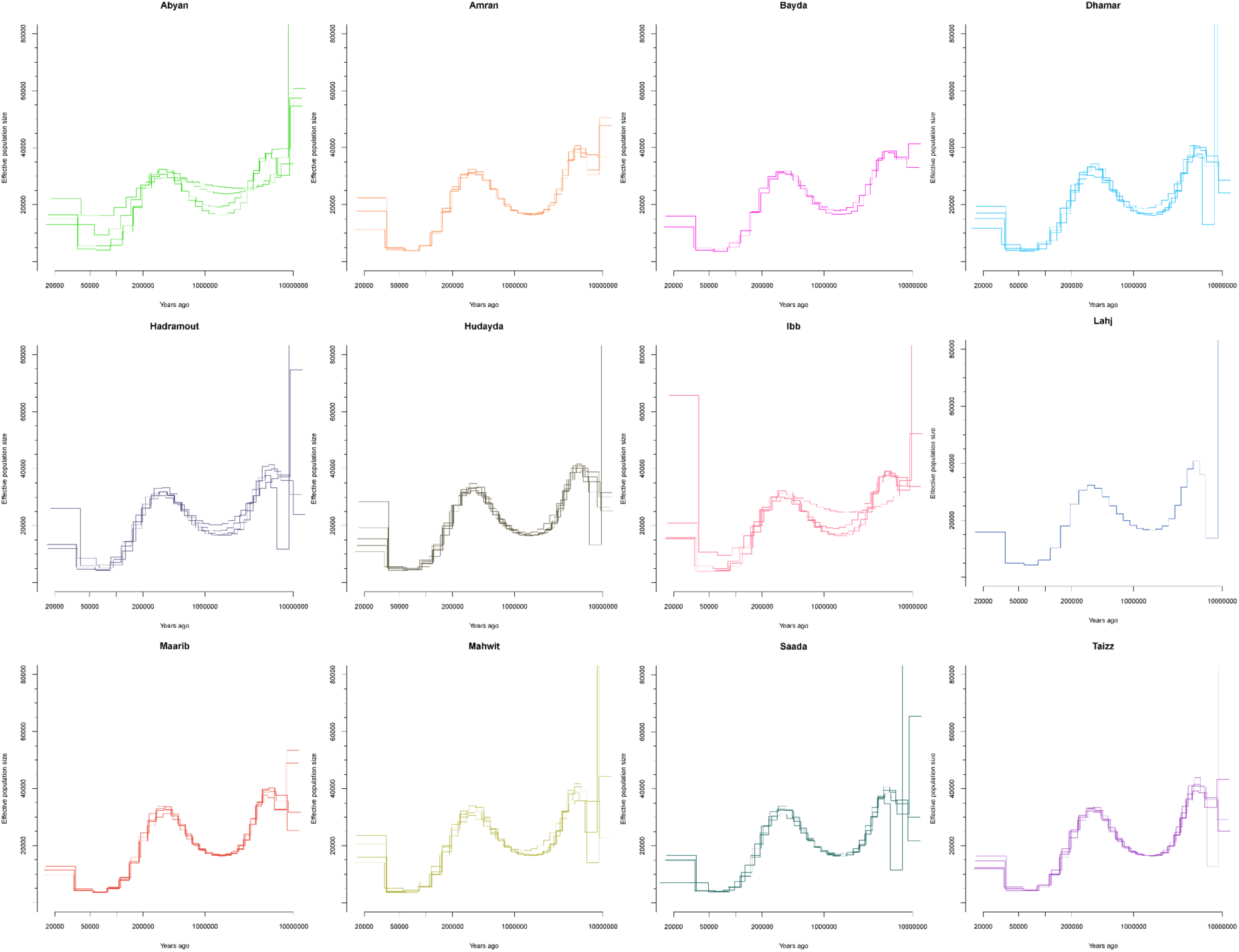
MSMC2 results showing effective population size change in Yemen over time. Most Yemeni genomes show a typical Eurasian population size change pattern characterized by a bottleneck around 50,000-60,000 years followed by a population size expansion.

**Figure 3.**
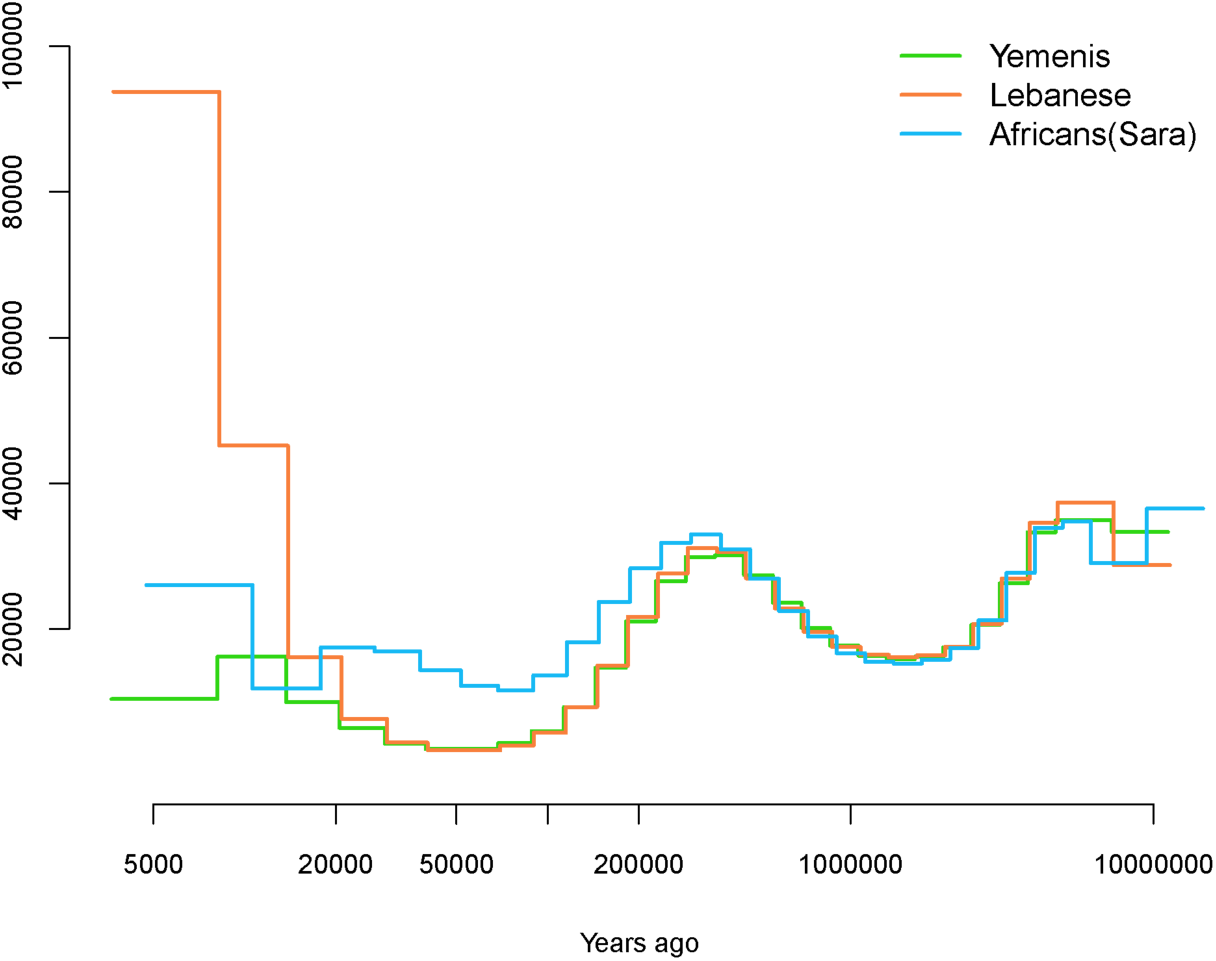
MSMC2 results showing population size change in Africans,^30^ Lebanese,^4^ and Yemenis using two genomes from each population.

**Figure 4.**
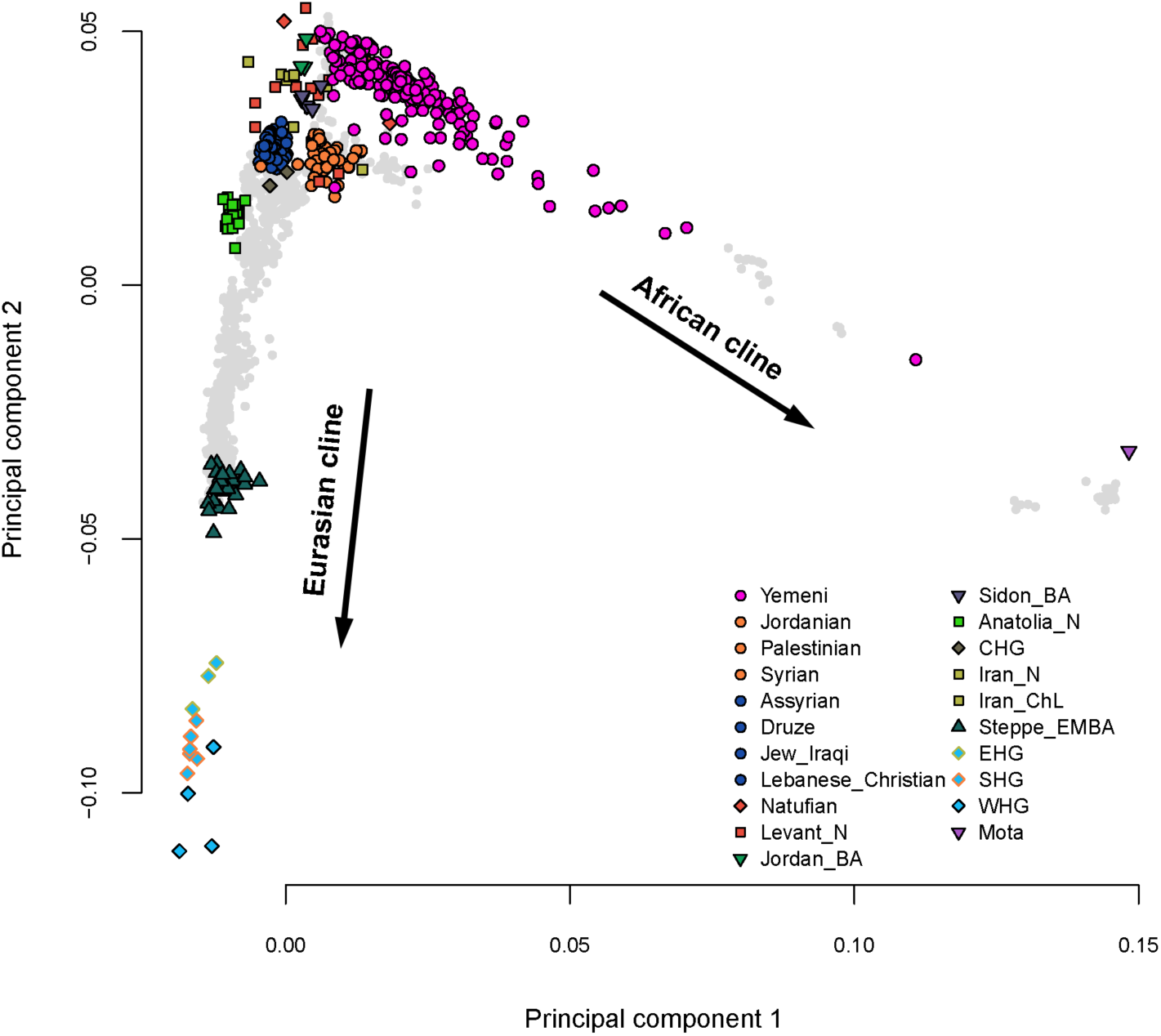
PCA with the Yemenis (pink circles) and the ancient samples projected into the plot using the variation from East Africans and West Eurasians (grey circles) including the Northern Near East populations (blue and orange circles).

**Figure 5.**
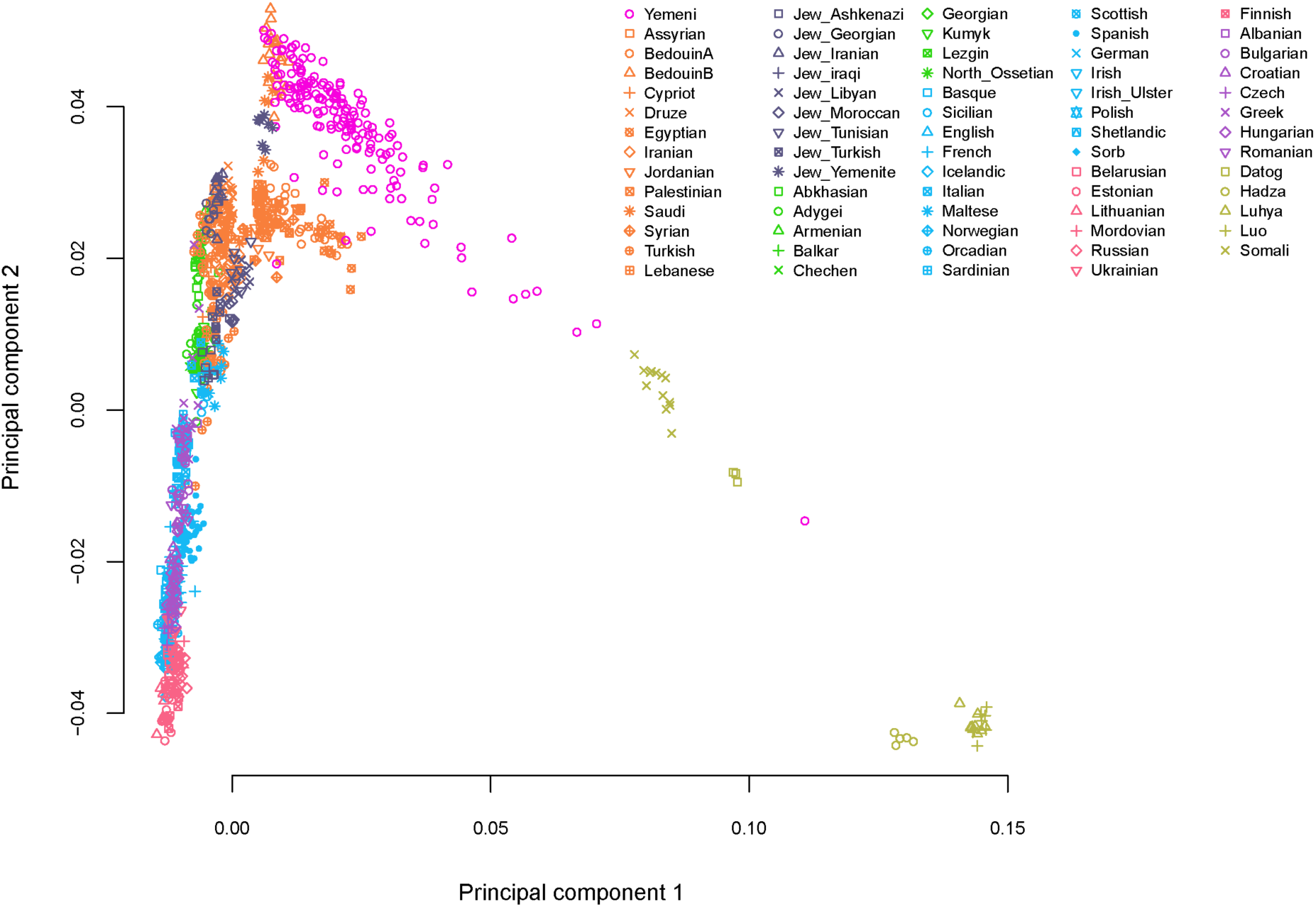
PCA as in Figure 4 but showing only the present-day populations.

**Figure 6.**
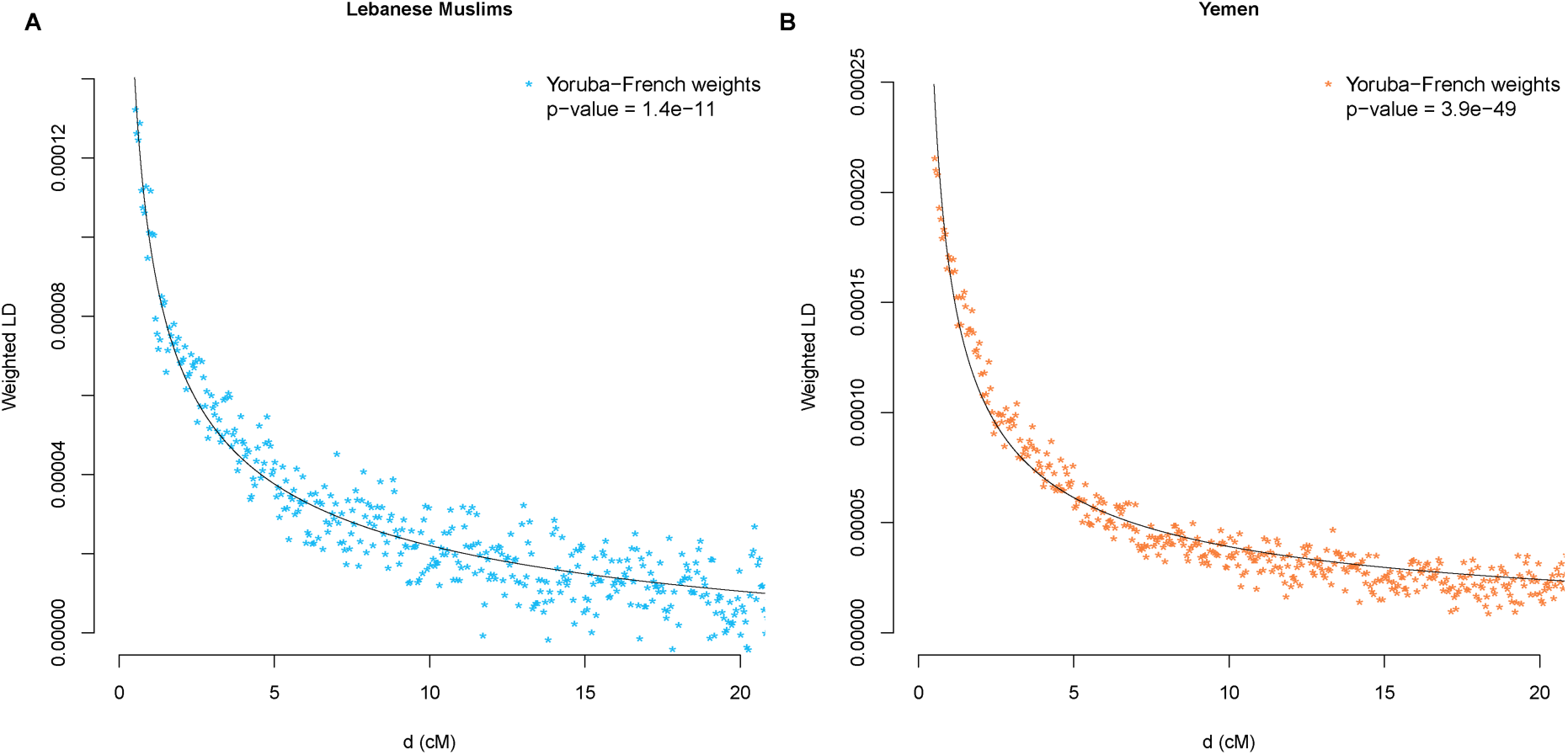
Weighted LD curves using Yoruba and French as reference populations for (A) Lebanese Muslims and (B) Yemenis.

Next, we investigated genetic structure within Yemen and found it was loosely correlated with geography (Figure 7): some clustering by region is visible, but many individuals from the same region do not cluster together. We interpret this structure as resulting from continuous movement of people between the different regions. However, we note that individuals from Hudayda and from Hadramout have increased African ancestry compared with most other Yemenis, while individuals from Maarib appear to be the least admixed clustering at the base of the African cline in Yemen. We confirm these results by running ADMIXTURE^20^ on the Yemenis, Lebanese, Africans and East Asians found in the *seq* set using K=3 to account for the African, Asian and Near Eastern ancestry. The ADMIXTURE results (Figure 8) confirm the observation from the PCA showing that most Yemenis have some African ancestry which is maximized in Hudayda and Hadramout, with the later having in addition some East Asian ancestry, while the Maarib individuals appear to be the least admixed of the populations tested in Yemen. The African and Asian ancestry in Yemen can also be observed in the present Y-chromosome and mtDNA lineages (Figure 9 and Table 3) which are enriched with Y-haplogroups A, E, and R1a and mtDNA haplgroups L and M, respectively.

**Table 3.** Y-chromosome and mtDNA haplogroups in Yemen

| ID | Y tree label | Region | Y haplogroup | MT Haplogroup |
| --- | --- | --- | --- | --- |
| EGAN00001456674 | Yemen-31 | Abyan | J-P58 | K1a27 |
| EGAN00001456628 | Yemen-45 | Abyan | R-Z92 | L0a2c |
| EGAN00001456614 | Yemen-20 | Abyan | J-P58 | M30+16234 |
| EGAN00001456688 | Yemen-42 | Abyan | J-M67 | U8b1a1 |
| EGAN00001456593 | Yemen-10 | Amran | G-F1761 | J1b |
| EGAN00001456579 | Yemen-16 | Amran | J-P58 | K1a7 |
| EGAN00001456663 | Yemen-39 | Amran | J-L27 | L3f1b+16292 |
| EGAN00001456648 | Yemen-6 | Amran | E-CTS1096 | M1a1f |
| EGAN00001456617 | Yemen-22 | Bayda | J-P58 | H2a1 |
| EGAN00001456603 | Yemen-5 | Bayda | E-CTS1096 | L2a1j |
| EGAN00001456575 | Yemen-15 | Bayda | J-P58 | M32c |
| EGAN00001456564 | Yemen-4 | Dhamar | E-CTS1096 | J1b2 |
| EGAN00001456684 | Yemen-34 | Dhamar | J-P58 | M1a1b1b |
| EGAN00001456668 | Yemen-30 | Dhamar | J-P58 | M1a1f |
| EGAN00001455435 | Yemen-3 | Dhamar | E-CTS1096 | U6a+16189 |
| EGAN00001456587 | Yemen-46 | Hadramout | R-Z2123 | J2a2a1 |
| EGAN00001456602 | Yemen-18 | Hadramout | J-P58 | L2a1b1a |
| EGAN00001456613 | Yemen-19 | Hadramout | J-P58 | M45a |
| EGAN00001456629 | Yemen-25 | Hadramout | J-P58 | U6a2a1 |
| EGAN00001456769 | Yemen-43 | Hudayda | J-M67 | J1b2 |
| EGAN00001456774 | Yemen-12 | Hudayda | J-CTS5368 | J1d1a1 |
| EGAN00001456561 | Yemen-44 | Hudayda | J-M205 | L2a1 |
| EGAN00001456745 | Yemen-8 | Hudayda | E-CTS1096 | L2a1c+16129 |
| EGAN00001456765 | Yemen-11 | Hudayda | J-CTS5368 | L2a1c+16129 |
| EGAN00001456568 | Yemen-1 | Ibb | A1b1-M118 | K2c |
| EGAN00001456592 | Yemen-17 | Ibb | J-P58 | L0a1a1 |
| EGAN00001456560 | Yemen-13 | Ibb | J-P58 | L0a1b1a1 |
| EGAN00001456637 | Yemen-27 | Ibb | J-P58 | U3b3 |
| EGAN00001456657 | Yemen-28 | Lahij | J-P58 | J1d1a |
| EGAN00001456627 | Yemen-2 | Lahij | E-L677 | J1d1a1 |
| EGAN00001456690 | Yemen-35 | Maarib | J-P58 | J1b2 |
| EGAN00001456634 | Yemen-26 | Maarib | J-P58 | J1d1a1 |
| EGAN00001456620 | Yemen-23 | Maarib | J-P58 | J2a2b |
| EGAN00001456675 | Yemen-32 | Maarib | J-P58 | U9a |
| EGAN00001456573 | Yemen-14 | Mahwit | J-P58 | HV1b1 |
| EGAN00001456650 | Yemen-7 | Mahwit | E-CTS1096 | L2a1a2 |
| EGAN00001456622 | Yemen-24 | Mahwit | J-P58 | L5a1a |
| EGAN00001456680 | Yemen-33 | Mahwit | J-P58 | T1a |
| EGAN00001456660 | Yemen-29 | Saada | J-P58 | HV1a'b'c |
| EGAN00001456615 | Yemen-21 | Saada | J-P58 | J1b1b1 |
| EGAN00001456705 | Yemen-41 | Saada | J-M47 | L2b1a3 |
| EGAN00001456676 | Yemen-40 | Saada | J-M47 | T1a2 |
| EGAN00001456743 | Yemen-38 | Taizz | J-P58 | H20 |
| EGAN00001456732 | Yemen-36 | Taizz | J-P58 | J2a2 |
| EGAN00001456737 | Yemen-37 | Taizz | J-P58 | L0a1a1 |
| EGAN00001456748 | Yemen-9 | Taizz | E-CTS1096 | R0a1a |

**Figure 7.**
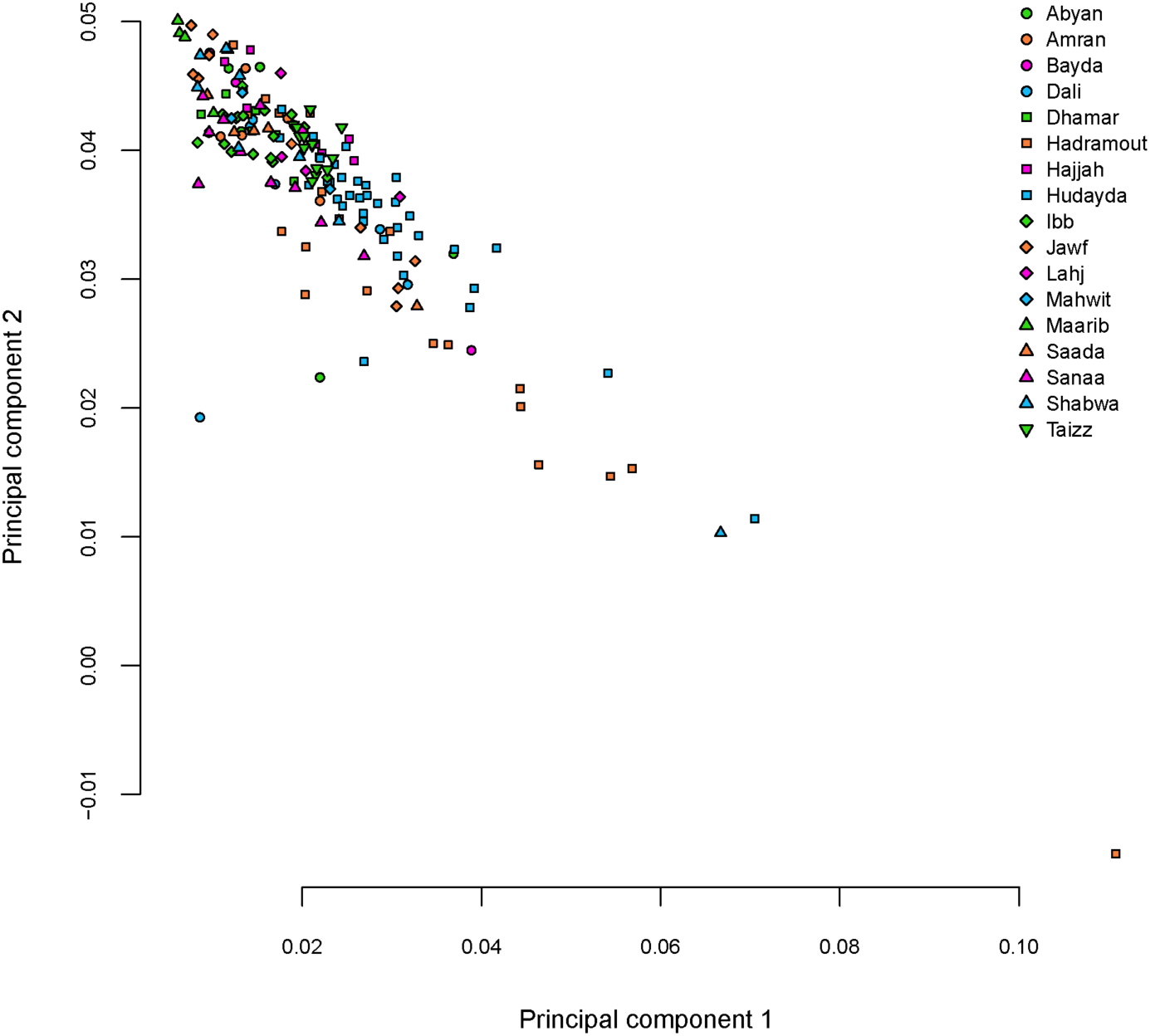
PCA coordinates from Figure 4 showing only samples from Yemen and their region of origin.

**Figure 8.**
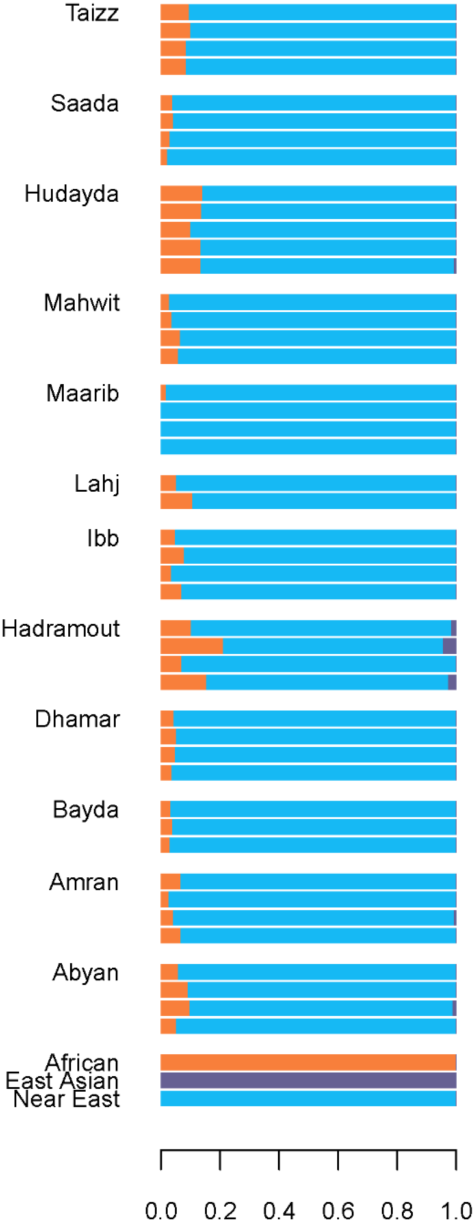
ADMIXTURE plot using the *geno* set and showing the Yemenis at K=3.

**Figure 9.**
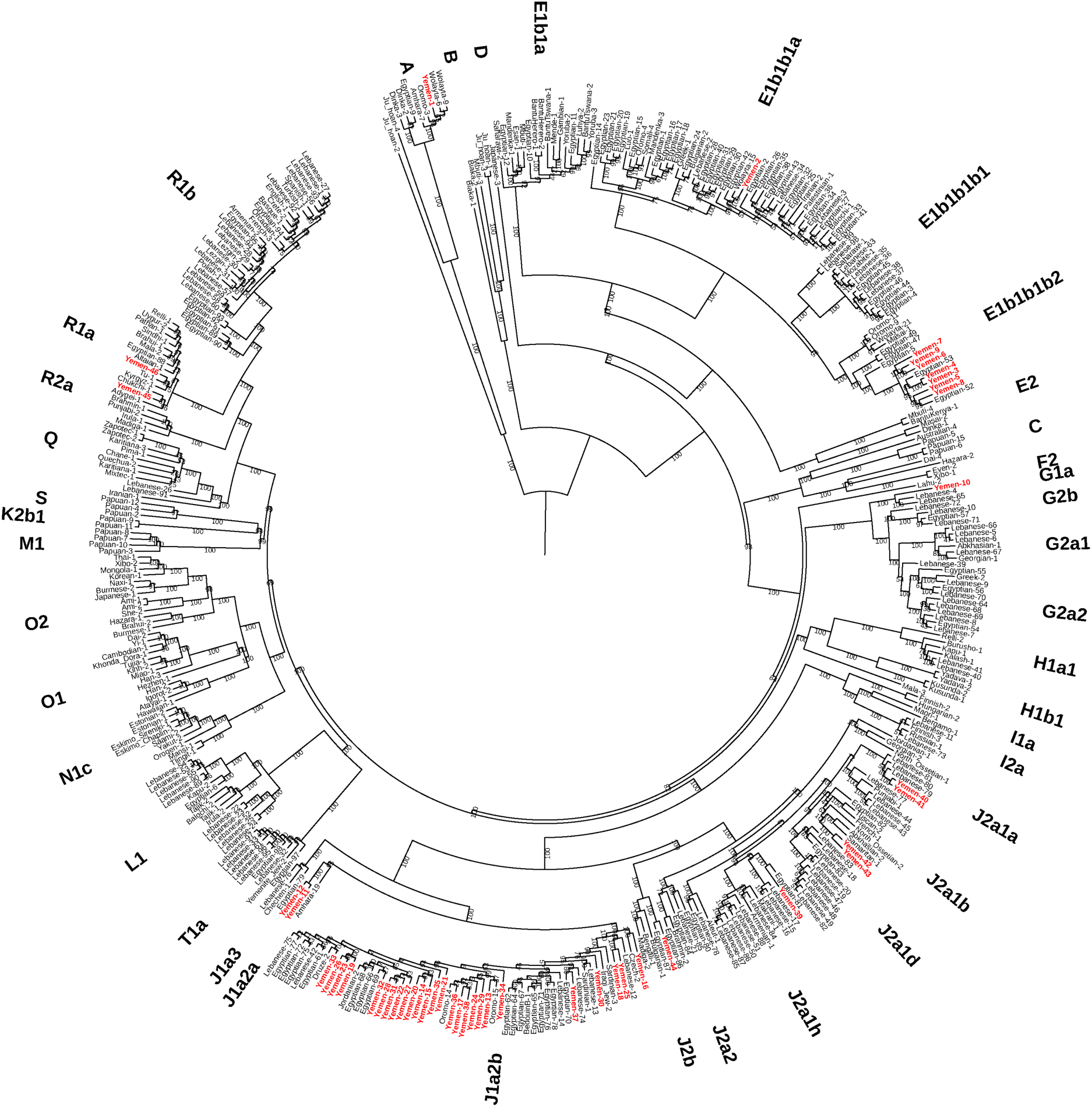
A maximum-likelihood tree of Y-Chromosome sequences from Yemen (red) and worldwide populations. Numbers on branches show bootstrap values from 1000 replicates.

The African ancestry in Yemen increases the genetic distance between the Yemenis and all other Eurasians and therefore prevents us from understanding how the Yemen’s Near Eastern ancestral component is related to other people in the region. But the relatively isolated Maarib individuals provide an opportunity to test Yemeni without the impact of the African ancestry. For example, using the four-population admixture test in ADMIXTOOLS,^9^ we find *f4*(Lebanese, Yemen; Ancient Eurasian, Chimp) is always positive with all ancient Eurasians being closer to Lebanese than to Yemenis (Figure 10A). But these results are biased by Yemen’s African ancestry since *f4*(Lebanese, Maarib; Ancient Eurasian, Chimp) shows Lebanese and Maarib are equally distant from the Neolithic (Z= 0.57) and Bronze Age (Z= 1.677) Levantine populations (Figure 10B). However, we detect Eurasian gene flow to the North but not to the South of the Near East, reflected by *f4*(Lebanese, Maarib; Steppe_EMBA, Chimp) being significantly positive (Z=10.837). We also note that Maarib has increased Natufian (Epipaleolithic hunter-gatherers) ancestry compared with the Lebanese, with *f4*(Lebanese, Maarib; Natufian, Chimp) being negative (Z= −2.95) (Figure 10B). We propose that the increase in Natufian ancestry in Maarib is likely from structure in the ancestral population. We confirm the *f4*-statistics results by modelling the modern and ancient Near Easterners using qpGraph from the ADMIXTOOLS package and show that our data support a model where the North and South of the Near East split from a population related to the Bronze Age population which inhabited the North of the Near East, but with the South regions having an excess of Epipaleolithic hunter-gatherer ancestry (Figure 11). After the split, the Northern regions of the Near East received gene flow from a Eurasian population^4; 19^ which never reached the South, where people instead admixed with Africans resulting in the genetic structure we observe in the Near East today (Figure 11).

**Figure 10.**
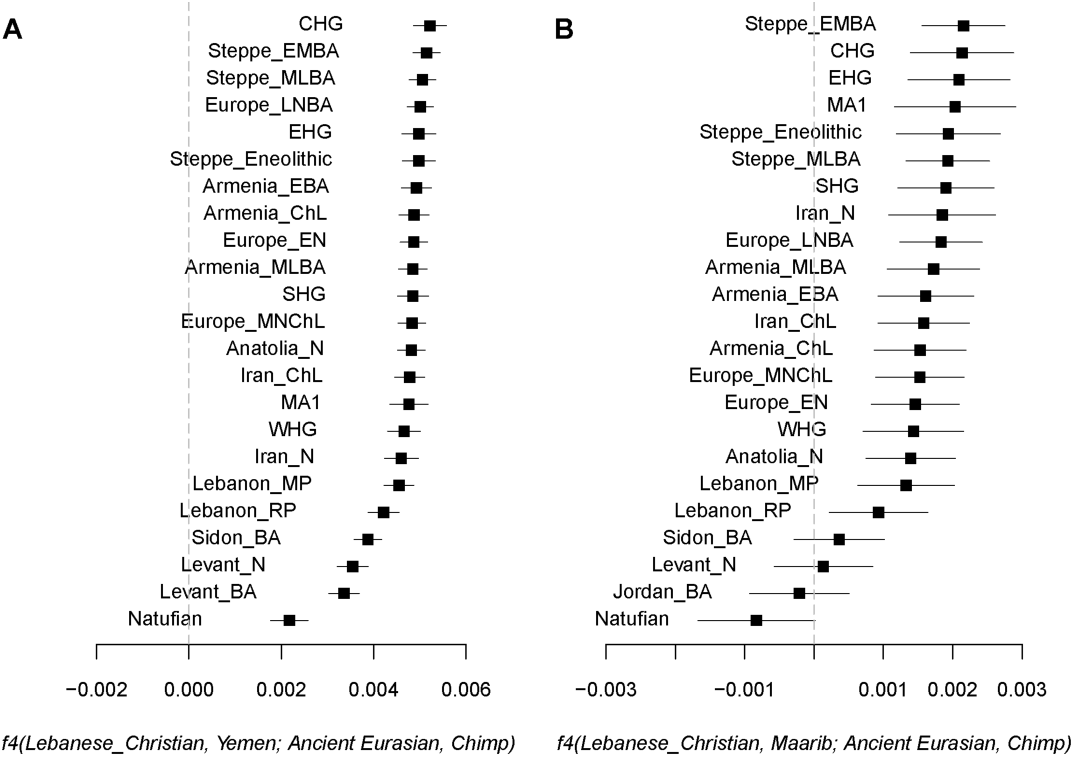
(A) The affinity of Yemen to ancient populations is masked by their African ancestry (B) but the less admixed Maarib population reveals the Near Eastern component in Yemen. We plot the *f4* statistic value and ±3 standard errors.

**Figure 11.**
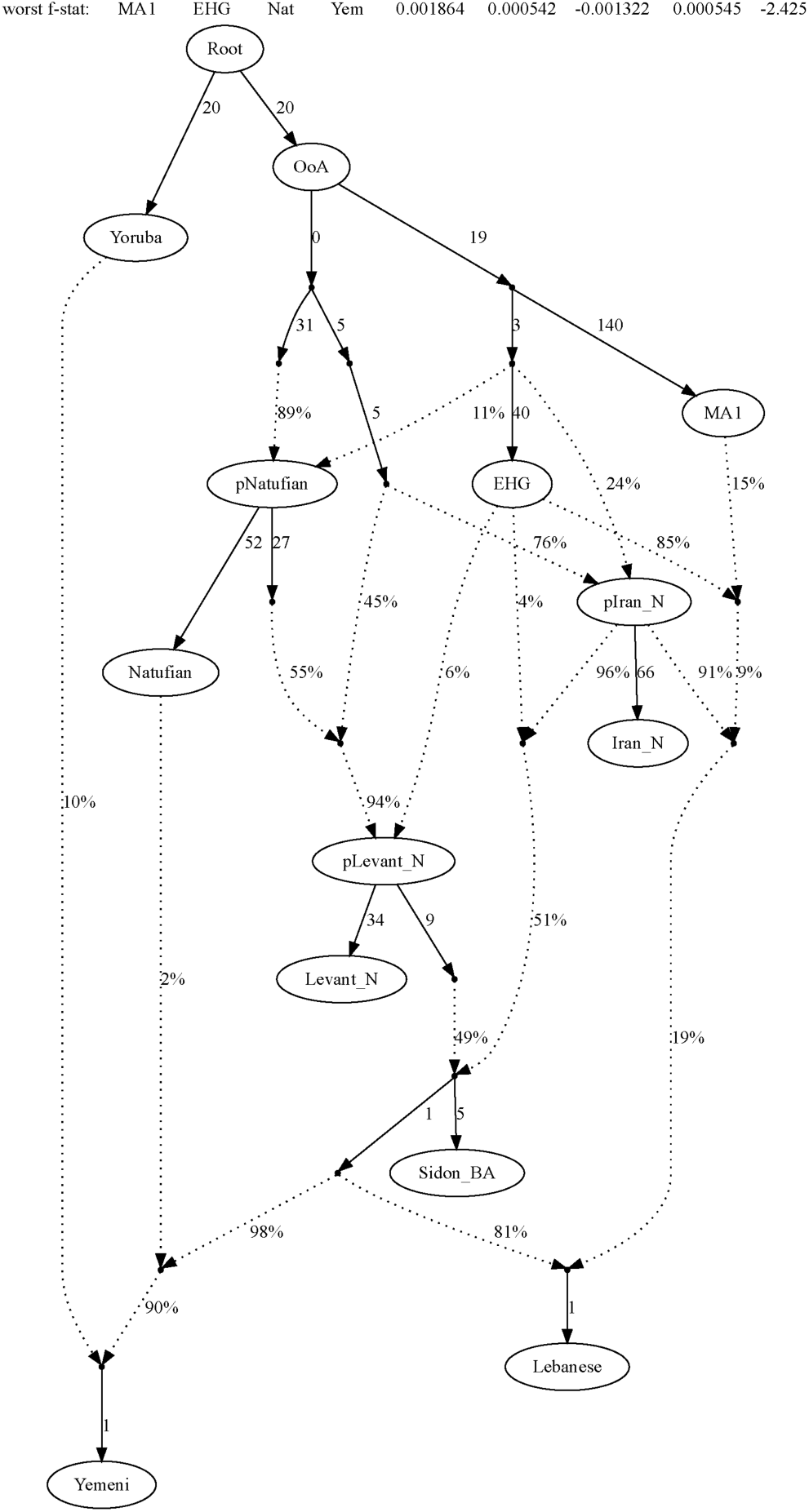
An admixture graph model showing the relationship of present-day Northern (Lebanese) and Southern (Yemeni) Near Easterners.

We thus find that the genetic structure of present-day Yemenis can be accounted for by mixture of post-glacial Levantine and other Eurasian populations with Africans within the last 1000 years. In particular, we do not find evidence for African ancestry that might have remained specifically in this region since the major expansion out of Africa 50,000-60,000 years ago.^1^ Our model of the genetic relationships in the Near East relies on the available genetic data today and thus will likely be refined in the future when, most importantly, ancient DNA data from the Southern regions of the Near East become available – work that has so far been hindered by the warm climate in the Near East negatively impacting the survival of ancient DNA.

## Methods

Array genotype data were processed using PLINK v1.90b5^21^ as described previously.^22^ Genotypes from sequence data were called using bcftools v1.6 with the command ‘bcftools mpileup -C50 -q30 -Q30 | bcftools call’ and phased with SHAPEIT v2.r900^23^ using the 1000 Genomes Project phase 3 haplotypes^2^ as a reference panel. Genotypes were merged with data from the literature using the program mergeit from the EIGENSOFT v7.2.1 package.^24^ Principal components were computed using smatpca v16000 also from the EIGENSOFT package with the following parameters; numoutlieriter: 0, lsqproject: YES, autoshrink: YES. Effective population size and rates of gene flow were inferred using MSMC v2.1.1^16^ with one (Figure 2) or two (Figure 3) high-coverage phased genomes from each population and using the following time segments -p 1*2+25*1+1*2+1*3, a generation time of 30 years and a rate of 1.25 × 10^-8^ mutations per nucleotide per generation. Y-chromosome haplogroups were assigned using yhaplo^25^ and phylogenetic analysis of the whole Y-chromosome sequences was carried out using RAxML v8.2.10^26^ with the arguments “-m ASC_GTRGAMMA” and “--asccorr=stamatakis” as described previously^6^ and plotted using iTOL v4.^27^ mtDNA haplogroups were determined using the mtDNA-server v1.0.7.^28^ Population mixture signals were tested using qpDstat v751 (f4mode: YES) and qpGraph v6412 (useallsnps: YES, blgsize: 0.05, forcezmode: YES, diag: .0001, bigiter: 6, hires: YES, lambdascale: 1, inbreed: YES) from the ADMIXTOOLS package.^9^ Admixture proportions were additionally estimated using ADMIXTURE v1.3.0.^29^ MALDER v1.0^17; 18^ (mindis: 0.005, binsize: 0.0005) was used to date the time of admixture using the Africans, East Asians and West Eurasians with n>20 in the *geno* set as references.

## Acknowledgements

Our work was supported by Wellcome grant number 098051 (MH, YC, YX, CTS). We thank Pierre Zalloua and Matthew Hurles for their contributions to this work.

## Data availability

Sequence data are available via EGAS00001002083, and genotype data via EGAS00001001231.

